# Tissue-specific heteroplasmy dynamics is accompanied by a sharp drop in mtDNA copy number during development

**DOI:** 10.1101/2021.07.30.454495

**Authors:** Nikita Tsyba, Maulik R. Patel

## Abstract

Mitochondrial mutation phenotypes are highly unpredictable as they depend on 3 variables; mutant-to-wildtype ratio (heteroplasmy level), total number of mitochondrial genomes (mtDNA), and the tissue affected. The exact phenotype experienced is governed by the combination of these variables, but current models lack the capability to examine the three variables simultaneously. We have established a *C. elegans* muscle and neuron system to overcome this challenge. Using this system, we measure heteroplasmy level and mtDNA copy number throughout development. Our results show that neurons accumulate significantly higher heteroplasmy level than muscles. These tissue-specific differences arise late in development, and are dependent on AMP-activated protein kinase (AMPK). Importantly, we find that somatic tissues lose more than half of their mtDNA content during development. These findings show that heteroplasmy levels can remain stable, or even increase, despite acute mtDNA losses.

## Introduction

The mitochondrial genome (mtDNA) is much smaller than the nuclear genome, but encodes core components of the electron transport chain (Gustafsson et al., 2016). Just like the nuclear genome, the mtDNA is susceptible to mutation that can disrupt function, leading to a wide range of seemingly unconnected symptoms (e.g. blindness, deafness, diabetes, cardiomyopathy) (Wallace and Chalkia, 2013). There are typically hundreds to thousands of mtDNAs in each cell (Robin and Wong, 1988, Veltri et al., 1990). Since every mtDNA replicates independently, multiple genotypes can arise within the same cell, leading to heteroplasmy (Pedreira et al., 2013). Some new genotypes are benign, while others may carry pathogenic mutations. Given the multi-copy nature of mtDNA, new mutations coexist with the functional genomes that can compensate for mutation effects (Shokolenko and Alexeyev, 2015). As a result, the emergence of pathogenic mutations does not immediately lead to metabolic disorders. However, as mutant mtDNAs proliferate and mutant-to-wildtype ratio rises, conditions become increasingly pathogenic. If the mutant mtDNA load rises beyond a critical threshold, the cell will not have sufficient wild-type mtDNA copies to meet energetic needs, leading to metabolic deficiency (Rossignol et al., 2003). However, mutant-to-wildtype ratio heteroplasmy level alone does not determine mutant phenotype.

In addition to heteroplasmy level, phenotypes depend on total mtDNA content and tissue type (Wallace and Chalkia, 2013, Filograna et al., 2019, Filograna et al., 2021, Li et al., 2015, Sharpley et al., 2012). Total mtDNA content represents the number of functional mitochondrial genomes available, and therefore a decrease in mtDNA content can have a profound effect on mitochondrial function. A decline in mtDNA copy number is demonstrated in mtDNA depletion syndromes (Filograna et al., 2021) and neurodegenerative disorders such as Parkinson’s and Alzheimer’s disease (Coskun et al., 2012, Rice et al., 2014). Copy number is also an important determinant of phenotype severity in heteroplasmic situations. For example, increasing mtDNA copy number by overexpressing mitochondrial transcription factor A alleviates phenotypic defects in heteroplasmic mice without altering mutant levels (Filograna et al., 2019, Jiang et al., 2017). Tissue type also affects mtDNA mutation phenotypes, with links to tissue energetic requirements (Wang et al., 2010). Cell types with high energetic requirements, like muscles and neurons, are especially sensitive to mitochondrial disorders (Wallace and Chalkia, 2013). As a result, muscular, cardiac and neurological dysfunction are among the most common symptoms caused by mitochondrial mutations (Wallace and Chalkia, 2013). Tissue-to-tissue differences in energetic requirements mean that the same heteroplasmy level may lead to a different phenotype depending on the tissue affected.

Differences in heteroplasmy level, mtDNA copy number and tissue physiology likely contribute to the heterogeneity of symptoms observed in patients (Wallace and Chalkia, 2013, Filograna et al., 2019, Filograna et al., 2021, Li et al., 2015, Sharpley et al., 2012, Rossignol et al., 2003). In fact, the exact phenotype experienced by a patient is presumably governed by the unique combination of these three variables. Consequently, taking all three parameters into account would help us better understand the phenotypic consequences of mtDNA mutations. Unfortunately, current models lack the capability to examine these three variables simultaneously, and the relationship between individual parameters as well as the mechanisms governing them remain largely unexplored. To address this problem, we established a *C. elegans* system that allows us to examine all three variables at the same time. *C. elegans* short lifespan, genetic tractability and mutant toolkit makes it a powerful system for mechanistic studies (Markaki and Tavernarakis, 2010, Corsi et al., 2015). Importantly, the *C. elegans* mtDNA is similar in size and gene content to that of mammals (Addo et al., 2010, Okimoto et al., 1992). It shares 12 of 13 protein coding genes, all 22 t-RNAs and two rRNAs with the mammalian genome. In addition, all *C. elegans* somatic cells are post-mitotic, which allows us to attribute differences in mutant load and copy number dynamics solely to intracellular mechanisms (Nussbaum-Krammer and Morimoto, 2014).

By capitalizing on these model advantages, we developed a method for measuring heteroplamy level and mtDNA copy number in tissue-specific manner. We use a combination of fluorescent-activated cell sorting (FACS) and droplet digital PCR (ddPCR) to isolate individual tissue types with single cell accuracy (Schmit et al., 2021). This level of precision ensures that we only collect cells from the tissue of interest and minimizes the risk of getting a mixed cell population. In addition, FACS allows us to keep the number of collected cells constant in each experiment. As a result, we can directly compare mtDNA content between experiments, which is crucial for mtDNA measurements. ddPCR provides the principal tool for measuring experimental mutant load and copy number by measuring the absolute number of DNA molecules and maintaining high precision at low template concentrations (Gitschlag et al., 2016, Hindson et al., 2011). In addition, ddPCR allows simultaneous measurement of both mutant and wildtype mtDNA levels, which makes heteroplasmy quantification more efficient. We applied these advantages to examine how heteroplasmy level and mtDNA content of *C. elegans* tissues change throughout development. We focused specifically on muscles and neurons, since these two cell types are especially sensitive to mitochondrial disorders (Wallace and Chalkia, 2013). Our results uncover tissue-to-tissue differences in heteroplasmy dynamics that were at least partially dependent on AMPK. The differences were accompanied by a sharp drop in somatic mtDNA content that continued throughout *C*.*elegans* development. Interestingly, the mtDNA decline did not lead to a decrease in heteroplasmy level. On the contrary, heteroplasmy level remained constant or even increased despite the mtDNA losses.

## Results

### Fluorescence activated cell sorting coupled with droplet digital PCR enables heteroplasmy measurement in specific cell types

Capitalizing on the existing collection of *C. elegans* strains with specific cell types fluorescently labeled we developed an approach to measure mutant frequency and copy number in a tissue-specific fashion. First, *C. elegans* cells are dissociated by dissolving the worm cuticle and connective tissues (Figure 1A). This method is highly scalable and enables generation of dissociated cell suspensions from a large population of animals. Next, the fluorophore-expressing cells are isolated with fluorescence activated cell sorting (FACS) (Figure 1A). FACS isolates individual cells that minimizes the risk of getting heterogeneous cell populations (Figure 1B and 1C) (Adan et al., 2017). FACS detects the fluorescence signal and light scatter parameters of individual cells and plots them on a 2-D chart. The 2-D chart allows us to draw a gate around cell populations of interest, which are then isolated by FACS (Figure 1C). To confirm that only viable cells are isolated, DAPI viability dye was added to each sample prior to sorting (Sauvat et al., 2015). Worms were age-synchronized by isolating embryos and growing them under identical temperature conditions. To maintain synchronization once worms become reproductively mature, experiments were performed with sterile animals using temperature-sensitive *glp-1(e2141)* strain. Homozygous *glp-1(e2141)* worms are sterile, if grown at 25°C restrictive temperature. Imaging experiments on FACS-collected neurons confirm the isolation of GFP-expressing cells (Figure 1D). We also show that isolated neuronal cells are viable and grow processes when maintained in cell culture (Figure 1E).

**Figure 1:**
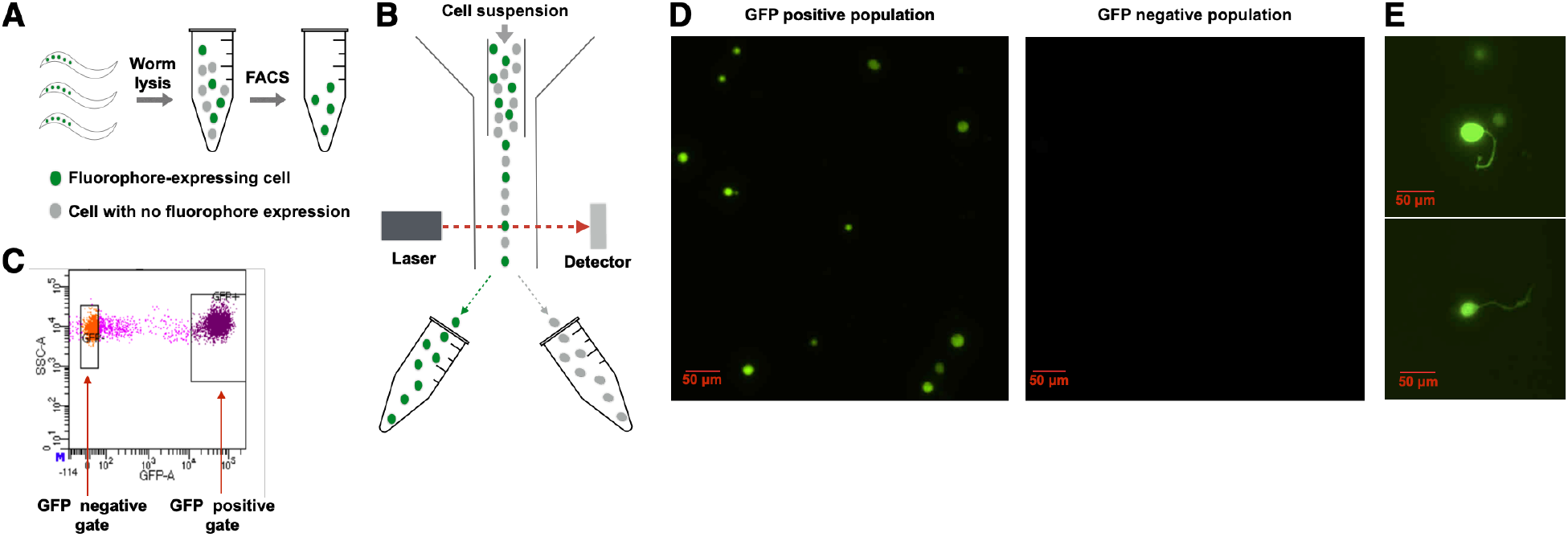
The design of the cell isolation protocol. [**A**] Graphical description of the cell isolation protocol: During lysis connective tissues are digested separating worm cells from each other. After that fluorophore-expressing cells are isolated with fluorescence-activated cell sorting (FACS). [**B**] Overview of the principle behind FACS-based cell isolation: First, cell suspension obtained after worm lysis enters fluidics system. The system divides cell suspension into droplets containing single cells. After that each cell is probed with a laser beam that measures the cell’s fluorescent signal. Finally, cells are sorted into separate compartments based on their fluorescent signal intensity. [**C**] FACS plot specifies which cell populations should be isolated, based on their GFP signal intensity. Each dot on the graph represents an individual cell. GFP signal intensity is on the x-axis, while side scatter parameter is on the y-axis. On this representative plot we specified GFP positive and GFP negative cell populations. FACS collects the two cell groups into separate compartments so that they can be used in subsequent experiments. [**D**] FACS enriches for GFP-expressing cells. The representative image on the left shows GFP-expressing cell enrichment in GFP positive population (corresponding to GFP positive gate in Figure 1C). The image on the right shows the lack of GFP positive cells in GFP negative population (corresponding to GFP negative gate in Figure 1C). The cells were cultured for 24 hours prior to imaging. [**E**] GFP positive neurons collected with FACS grow processes in cell culture. Representative images demonstrating that FACS collects the correct cell type and that the collected cells are viable after 24 hours in cell culture.

Once cells of interest are collected, we measure their heteroplasmy levels and copy number with droplet digital PCR (ddPCR). The advantages of this technology stem from sample partitioning prior to PCR reaction. In ddPCR, the sample is partitioned into thousands of droplets and PCR reactions occur independently in each individual droplet. Low sample concentrations ensure that each droplet encloses no more than 1 mtDNA copy (Hindson et al., 2011). Since droplets contain PCR product derived from a single mtDNA molecule, mutant and wild-type-carrying droplets can be identified based on their fluorescent signal (Figure 2A, 2B). As a result, mutant and wildtype mtDNA levels can be measured by counting the absolute number of droplets carrying either genome. We have successfully optimized the ddPCR protocol for two different mtDNA mutations–*uaDf5* and *mptDf2*. The two mutations are large non-overlapping deletions affecting multiple mtDNA genes (Figure 2C). Both deletions exist in a state of heteroplasmy with the wildtype genome. The primers we used allow us to reliably separate mutant and wildtype-carrying droplets for both heteroplasmy strains (Figure 2B).

**Figure 2:**
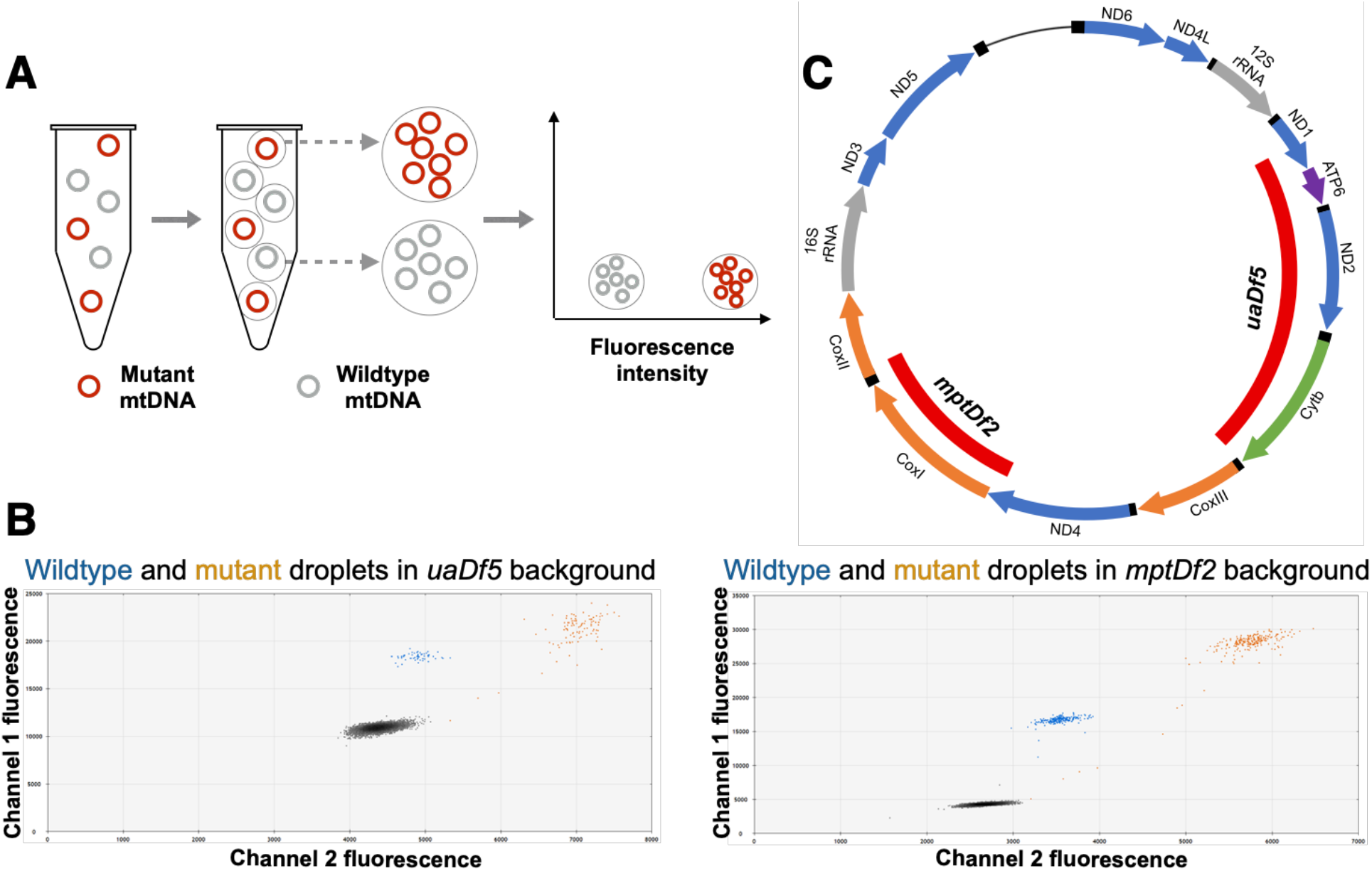
The principle behind ddPCR and its application for heteroplasmy level measurements. [**A**] Overview of the principle behind droplet digital PCR: First, a sample is partitioned into individual droplets. Low sample concentration ensures that each droplet contains at maximum a single mtDNA molecule. After partitioning independent PCR reactions occur in each individual droplet. Finally, mutant and wild-type-carrying droplets are identified based on their fluorescence intensity. [**B**] ddPCR fluorescence intensity plot showing the separation between mutant and wild-type-carrying droplets. The plots demonstrate droplets’ signal intensity in two fluorescent channels. Each dot represents an individual droplet carrying a single mtDNA molecule. Droplets carrying mutant mtDNA have higher fluorescence and are labeled with orange color. Wild-type-carrying droplets are labeled blue. The plots demonstrate the separation between mutant and wild-type-carrying droplets in *uaDf5* and *mptDf2* mutant backgrounds (left and right charts, respectively). [**C**] *C*.*elegans* mtDNA map showing the positions of two deletion mutations used in our experiments: *uaDf5* and *mptDf2*.

### Mutant load increases in neurons but not muscles during development

Muscles and neurons have some of the highest energy demands and are affected severely by mtDNA mutations (Wang et al., 2010, Wallace, 2005). Heteroplasmic mutant strains were crossed into transgenic lines expressing GFP pan-neuronally or in body wall muscles (Altun-Gultekin et al., 2001, Haynes et al., 2007). The best studied *C. elegans* mtDNA deletion (3.1 Kb *uaDf5*) (Tsang and Lemire, 2002b) affects multiple protein-coding genes and tRNAs (Figure 2B) to impact cellular respiration, increase mitochondrial stress and reduce viability, with lower brood size and slower growth rate (Liau et al., 2007, Lin et al., 2016, Gitschlag et al., 2020). We first examined *uaDf5* levels in neurons and muscles of adult animals (Day 1) from pools of 200 cells, which provided sufficient mtDNA templates for ddPCR while also allowing us to collect multiple batches of cells. To test whether mutant load in the GFP-positive cells differs from the rest of the tissues, we collected total somatic cells and used them as an internal control; mutant load in these cells represents the average *uaDf5* level in *C. elegans* soma.

There was no significant difference in the *uaDf5* levels between body wall muscle cells and total somatic cells in Day 1 adults (Figure 3A). In contrast, neurons had 7.6% higher *uaDf5* levels than total somatic cells (p-value = 0.0006, unpaired t-test) (Figure 3A). These results suggest that muscles and neurons have distinct heteroplasmy levels. However, the *uaDf5* levels in neurons and muscles cannot be compared directly because they were measured in separate worm populations. Therefore, we normalized the heteroplasmy level in each tissue to its corresponding total somatic cell control (Burgstaller et al., 2014, Johnston and Jones, 2016, Latorre-Pellicer et al., 2019). Comparing normalized values confirmed that neurons have higher *uaDf5* levels than muscles (Figure 3B). This difference may be the result of inheriting distinct mutant load during embryogenesis. Alternatively, the heteroplasmy levels may have diverged post-mitotically. To differentiate between these two possibilities, we repeated experiments in newly hatched L1 larvae that already have most neurons in place (Sulston, 1976, Sulston and Horvitz, 1977). We found no significant difference in *uaDf5* levels between L1 tissues and the corresponding total somatic cell controls (Figure 3C). These data suggest that while the mutant load remains static in muscles, it increases in neurons throughout larval development. We formally tested and confirmed this possibility by comparing normalized *uaDf5* levels between larval L1 and Day 1 adult animals (Figure 3D). These results uncover the differences in heteroplasmy dynamics between neurons and body wall muscles. Importantly, the differences manifest themselves at late developmental stages and are not present in the newly hatched animals, suggesting they are caused by post-mitotic mechanisms.

**Figure 3:**
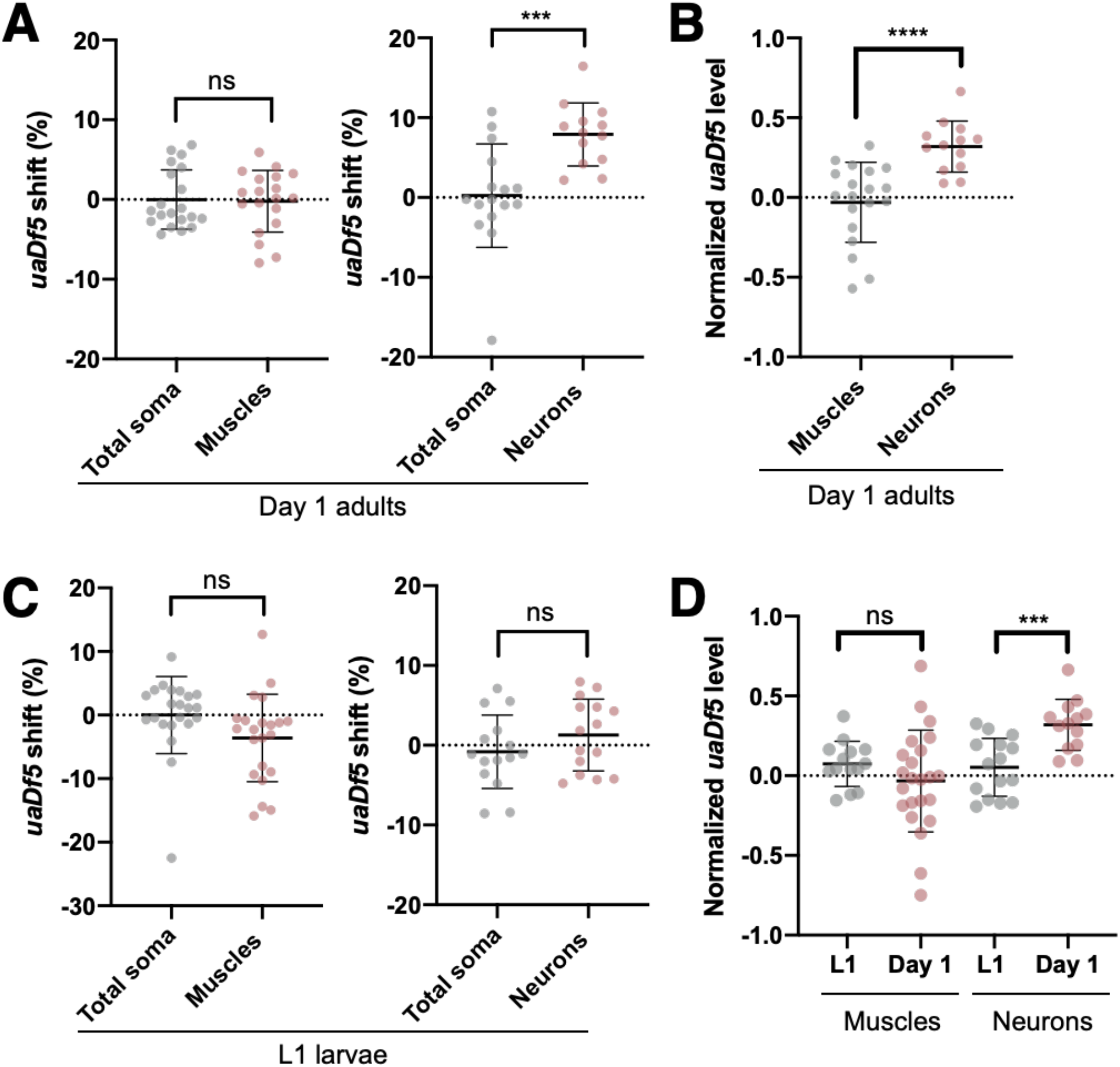
Neurons, but not muscles, accumulate higher *uaDf5* levels than the rest of the cells. [**A**] *uaDf5* level in muscles and neurons relative to cell suspension control at Day 1 of adulthood (N = 19 (20) for muscles, N = 13 (16) for neurons, N of the corresponding control groups is given in brackets). The control contains all viable cells from the lysed animals and is derived from the same cell population as the tissue of interest. Since each tissue was isolated from a separate worm population, muscles and neurons are compared to their own cell suspension controls (total soma). [**B**] Normalized *uaDf5* level in muscles and neurons of Day 1 adult animals (N = 17 for muscles, N = 20 for neurons). [**C**] *uaDf5* level in muscles and neurons of L1 larva relative to cell suspension control (N = 22 (22) for muscles, N = 15 (15) for neurons). [**D**] *uaDf5* shift between L1 and Day 1 time points in muscles and neurons (Muscles: N = 14 for Day 1, N = 23 for L1. Neurons: N = 13 for Day 1, N = 15 for L1). To enable comparisons between the time points *uaDf5* level of each sample in panels B and D was normalized to its corresponding total soma control. These panels represent the normalized version of the data from panels A and C, except for muscles data from panel D. Statistical tests: all comparisons in this figure were made using unpaired t-test with Welch’s correction.

### The cell-type specific mutant mtDNA dynamics are generalizable to the *mptDf2* deletion

To determine whether the results are specific to *uaDf5* or generalizable across heteroplasmies, we performed similar experiments with a different mtDNA deletion (*mptDf2*) (Thompson et al., 2013). Like *uaDf5, mptDf2* is a large mtDNA deletion involving multiple protein-coding genes and tRNAs. However, *mptDf2* and *uaDf5* affect distinct mtDNA regions and genes (Figure 2C). Despite these differences, experiments with *mptDf2* yielded similar results. Neurons accumulated 10.46% higher *mptDf2* levels than the rest of the somatic cells by the first day of adulthood (p-value = 0.0021, unpaired t-test) (Figure 4A). In contrast, muscles retained the same mutant load as total somatic cells (Figure 4A). Comparison of normalized values shows that neurons have significantly higher *mptDf2* level than muscles (Figure 4B). Experiments with L1 animals show no difference in *mptDf2* levels between the tissues and total somatic cells during early development (Figure 4C). This analysis confirms that mutant load increases in neurons but remains unchanged in muscles across development (Figure 4D). Collectively, these data indicate that *mptDf2* exhibits similar heteroplasmy dynamics to *uaDf5* despite the differences in genes affected. As with *uaDf5, mptDf2* level remained static in body wall muscles, while increasing significantly in neurons throughout *C*.*elegans* development. The heteroplasmy differences manifested themselves at late developmental stages indicating that the difference in *mptDf2* level is also mediated by post-mitotic mechanisms.

**Figure 4:**
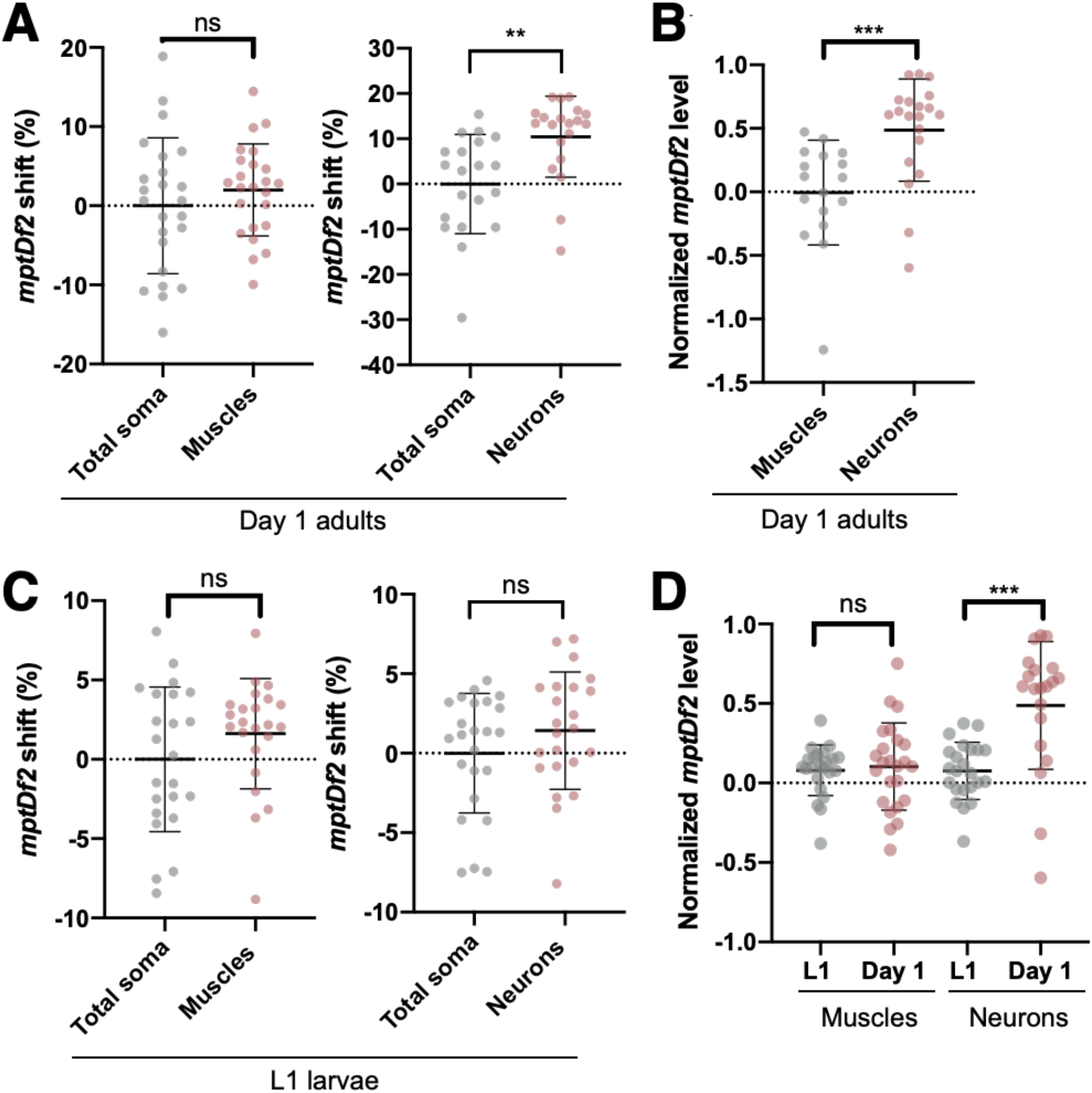
Neurons, but not muscles, accumulate higher *mptDf2* levels than the rest of the cells. [**A**] *mptDf2* levels in muscles and neurons relative to cell suspension control at Day 1 of adulthood (N = 24 (24) for muscles, N = 20 (20) for neurons, N of the corresponding control groups is given in brackets). [**B**] Normalized *mptDf2* level in muscles and neurons of Day 1 adult animals (N = 19 for muscles, N = 13 for neurons). [**C**] *mptDf2* level in muscles and neurons of L1 larva relative to cell suspension control (N = 23 (23) for muscles, N = 23 (24) for neurons, N of the corresponding control groups is given in brackets). [**D**] *mptDf2* shift between L1 and Day 1 time points in neurons and muscles (Neurons: N = 20 for Day 1, N = 23 for L1. Muscles: N = 23 for Day 1, N = 23 for L1). To enable comparisons between the time points *mptDf2* level of each sample in panels B and D was normalized to its corresponding cell suspension control. These panels represent the normalized version of the data from panels A and B, with the exception of muscles data from panel B. Statistical tests: all comparisons in this figure were made using unpaired t-test with Welch’s correction.

### AMPK modulates mutant load in muscles but not in neurons

We next sought to identify the mechanisms that modulate tissue-specific heteroplasmy dynamics. Since tissue-to-tissue differences are not present in L1 larvae, they could not be caused by biased mutant mtDNA inheritance. Furthermore, since there is no cell turnover in *C. elegans*, heteroplasmy differences can only arise through intracellular mechanisms (Rajasimha et al., 2008). The energy sensor AMP-activated protein kinase (AMPK) provides one such mechanism as it regulates the equilibrium between mitochondrial biogenesis and degradation (Palikaras and Tavernarakis, 2014, Herzig and Shaw, 2018). Importantly, AMPK is differentially expressed among *C. elegans* tissues (Lee et al., 2008). We therefore hypothesized that AMPK affects heteroplasmy level in a tissue-specific fashion and contributes to the mutant load differences between muscles and neurons.

To test this hypothesis, we examined the impact of AMPK on *uaDf5* levels in muscles and neurons by knocking out the alpha subunit of AMPK (*aak-2)* (Apfeld et al., 2004). In body wall muscles, AMPK knockout led to a significant increase in *uaDf5* levels, but only in Day 1 adult animals (p-value <0.0001, unpaired t-test) (Figure 5A). The *uaDf5* level of L1 muscles remained similar that of total somatic cells despite the knockout (Figure 5A). This result suggests that the effects of AMPK knockout manifest themselves at later developmental stages. Comparison of normalized *uaDf5* levels between L1 and Day 1 animals confirms this suggestion by showing a significant heteroplasmy increase across development (Figure 5B). This increase in muscles’ *uaDf5* level does not occur in wild-type animals (Figure 3D). Consistent with these data, the *aak-2* mutants have higher normalized *uaDf5* levels than their wild-type counterparts (p-value = 0.0003, unpaired t-test) (Figure 5C). These results show that AMPK knockout leads to a significant increase in heteroplasmy level in body wall muscles. Importantly, the increase occurred at later developmental stages suggesting that AMPK affects mutant load through post-mitotic mechanisms.

**Figure 5:**
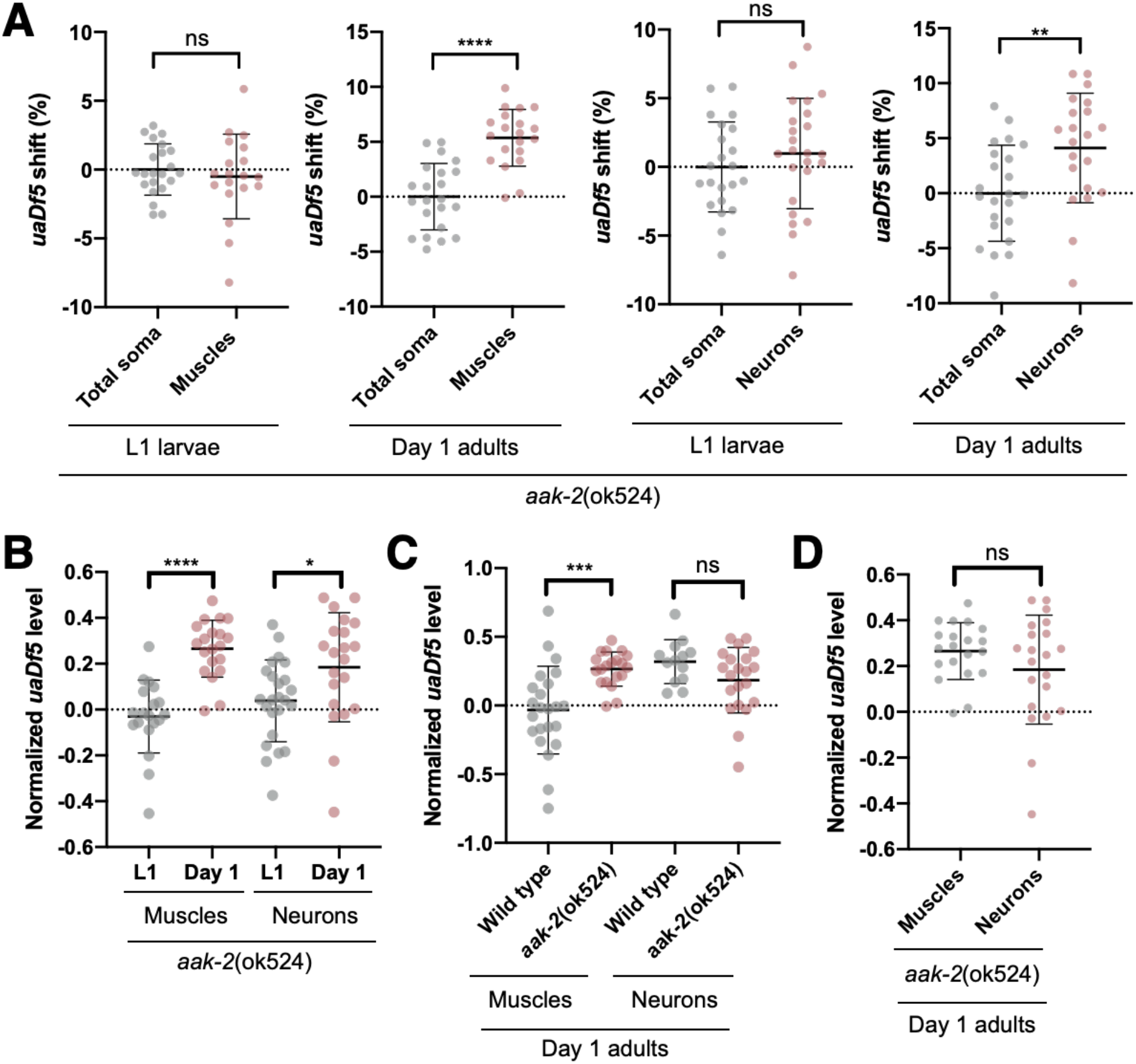
AMPK knockout causes *uaDf5* levels to increase in muscles, but not in neurons. [**A**] *uaDf5* shift in muscles and neurons of *aak-2 (ok524)*, AMPK knockout animals relative to cell suspension control at L1 and Day 1 time points (N = 20 (22) for Day 1 muscles, N = 19 (22) for L1 muscles, N = 21 (23) for Day 1 neurons, N = 24 (22) for L1 neurons, N of the corresponding control groups is given in brackets). [**B**] *uaDf5* shift in *aak-2 (ok524)* mutant muscles and neurons between L1 and Day 1 time points (Muscles: N = 20 for Day 1, N = 19 for L1. Neurons: N = 21 for Day 1, N = 24 for L1). [**C**] The effect of AMPK knockout on *uaDf5* level in muscles and neurons (Muscles: N = 20 for AMP-/-, N = 23 for wild-type. Neurons: N = 21 for AMP-/-, N = 13 for wildtype). [**D**] The difference in *uaDf5* level between muscles and neurons disappears after AMPK knockout (N = 20 for muscles, N = 21 for neurons). To enable comparisons between the time points *uaDf5* level of each sample in panels B, C and D was normalized to its corresponding cell suspension control. These panels represent the normalized version of the data from figures 5A and 3C. Statistical tests: all comparisons in this figure were made using unpaired t-test with Welch’s correction.

In contrast to muscles, the loss of AMPK does not significantly affect *uaDf5* mutant load in neurons. Although there was no significant difference in *uaDf5* levels between neurons and total somatic cells at the L1 stage, by Day 1 adulthood neurons accumulated 4.1% higher *uaDf5* levels (p-value = 0.0059, unpaired t-test) (Figure 5A). However, this increase in *uaDf5* level was already present in the wildtype background. The comparison of normalized *uaDf5* levels shows no significant difference between AMPK mutant and wild-type neurons (Figure 5C). The data show that AMPK knockout does not have a significant effect on heteroplasmy in neurons. This finding is in contrast with our observations from body wall muscles, which were significantly affected by the knockout. Overall, our results indicate that the loss of AMPK differentially affects heteroplasmy level in muscles and neurons. In turn, it suggests that AMPK functions in a tissue-specific fashion to modulate heteroplasmy levels. Consistent with this idea, the *uaDf5* level difference between neurons and muscles disappears in the absence of AMPK (Figure 5D compared to Figure 3B). In summary, these data suggest that AMPK contributes to heteroplasmy differences between body wall muscles and neurons.

### Somatic mtDNA levels decrease dramatically during development and aging

In addition to mutant load, changes in total mtDNA copy number can also contribute to pathogenicity, affecting the number of wild-type functional mtDNA copies (Filograna et al., 2021). The same heteroplasmy level may lead to different phenotypes depending on the mtDNA content of a cell, which makes copy number an important modulator of mtDNA mutations’ symptoms severity (Figure 6A) (Filograna et al., 2019). Therefore, we measured total mtDNA content in muscles and neurons at both larval L1 and adult Day 1 time points. Both tissues exhibit a sharp drop in mtDNA levels within this period, losing more than 70% of their mtDNA content (Figure 6B). We observed a similar dramatic decrease in total somatic cells, suggesting that this drop is not confined to neurons and muscles (Figure 6C). Furthermore, the data from Day 8 adults show that mtDNA levels continue to decline as worms age (Figure 6D). Somatic cells lose more than 80% of their starting mtDNA content by day 8 of adulthood. Finally, to determine whether the decrease in mtDNA copy number is generalizable, we analyzed the mtDNA levels in heteroplasmic animals harboring the *mptDf2* deletion. We observed a similarly sharp drop in mtDNA levels even in *mptDf2* background, where muscles, neurons and total somatic cells lost more than 50% of their mtDNA by the first day of adulthood (Figure 7A-B). Taken together, these data suggest that the decline in mtDNA content is likely a general feature of somatic cells in *C. elegans*.

**Figure 6:**
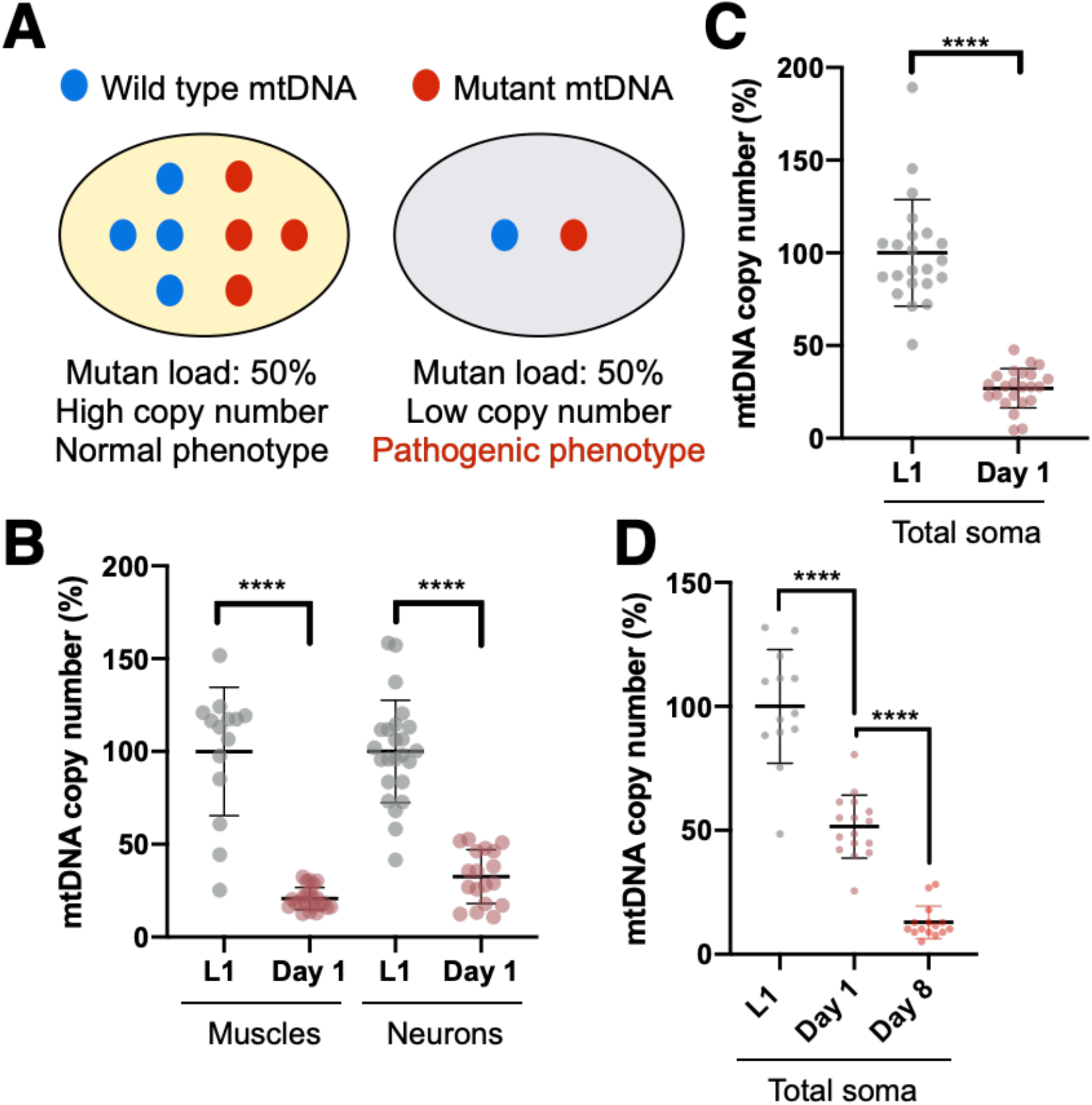
mtDNA content decreases sharply between L1 and Day 1 adulthood time points. The experiments presented in this figure have been carried out in *uaDf5* background. **[A**] Total mtDNA copy number can modulate the phenotype of mtDNA mutations. [**B**] mtDNA content drops sharply in both muscles and neurons between L1 and Day 1 time points (Muscles: N = 23 for Day 1, N = 14 for L1. Neurons: N = 18 for Day 1, N = 24 for L1). [**C**] The drop in mtDNA copy number is not restricted to muscles and neurons. It occurs in all somatic cells between L1 and Day 1 time points (N = 23 for Day 1, N = 22 for L1). [**D**] Somatic mtDNA content continues to decrease after the first day of adulthood (N = 16 for Day 1, N = 13 for L1, N = 15 for Day 8). Statistical tests: all comparisons in this figure were made using unpaired t-test with Welch’s correction.

**Figure 7:**
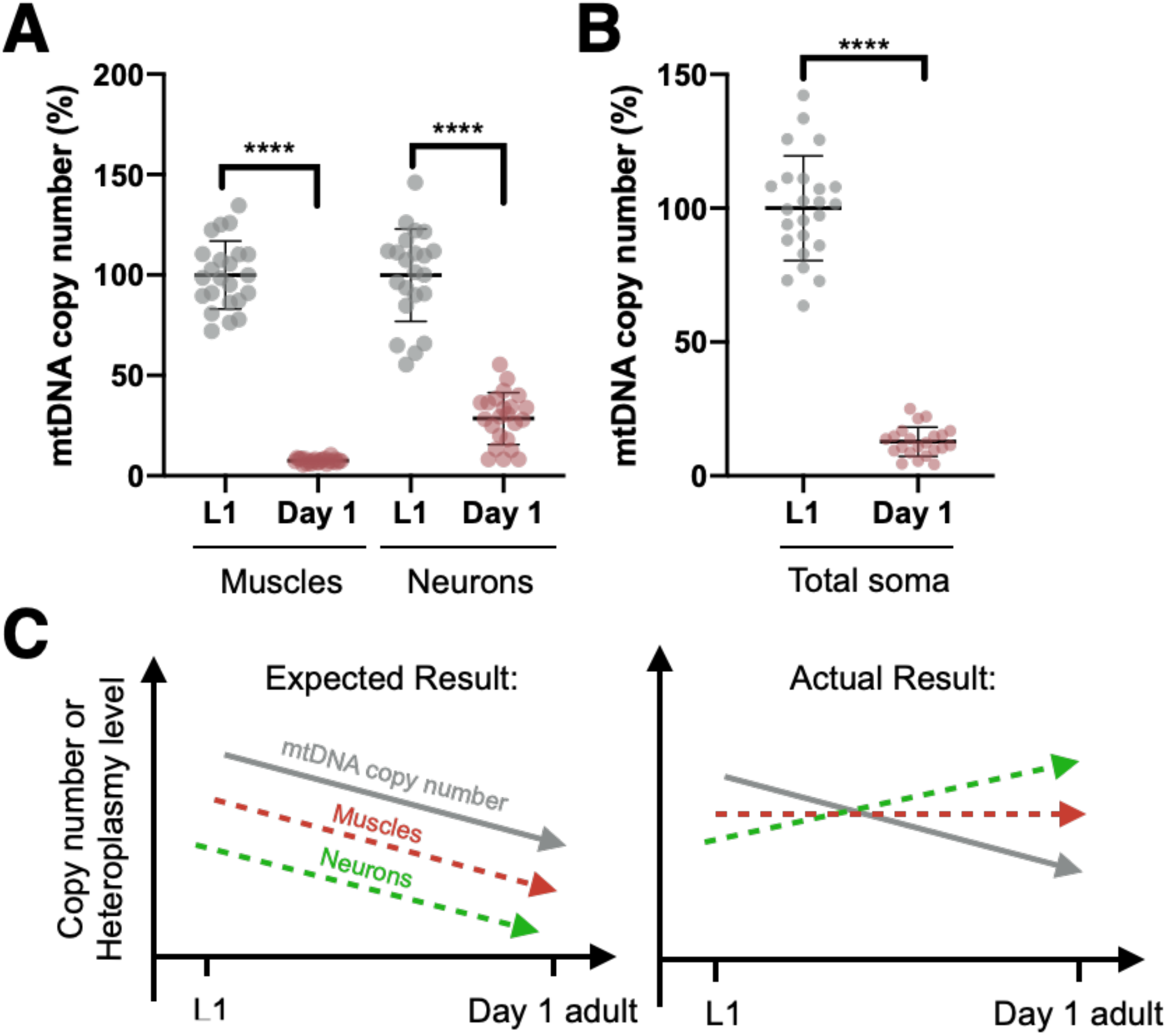
The sharp drop in mtDNA content persists in *mptDf2* mutation background. [**A**] Change in mtDNA levels in muscles and neurons between L1 and Day 1 time points (Muscles: N = 24 for Day 1, N = 23 for L1. Neurons: N = 23 for Day 1, N = 22 for L1). [**B**] Change in mtDNA content of all somatic cells (N = 22 for Day 1, N = 24 for L1). [**C**] In contrast to the expected outcome, the sharp drop in mtDNA copy number (solid grey line) did not lead to a comparable decrease in heteroplasmy levels (colored dashed lines). Statistical tests: all comparisons in this figure were made using unpaired t-test with Welch’s correction.

## Discussion

Extreme phenotypic diversity is a prominent feature of mtDNA mutations (Wallace, 2005, Wallace and Chalkia, 2013, Rossignol et al., 2003). A given mutation may cause severe symptoms in one patient, such as blindness and deafness, but lead to no symptoms in another (Rossignol et al., 2003). Understanding the factors that determine this variation is of utmost importance for mitochondrial research. Previous studies indicate that the mutation phenotype likely depends on heteroplasmy level, mtDNA content and tissue type (Filograna et al., 2019, Rossignol et al., 2003, Wallace, 2005). In this study, we developed a *C. elegans* model to measure heteroplasmy level and copy number in a tissue-specific fashion. Our results reveal significant differences in heteroplasmy between muscles and neurons. We found that adult neurons, but not muscles, accumulate significantly higher heteroplasmy levels. This finding has several important implications. First, it suggests that muscles and neurons have distinct heteroplasmy dynamics, despite being similar energetically demanding tissues. Second, it rules out unequal mutant mtDNA inheritance as a possible mechanism causing the difference, suggesting a post-mitotic mechanism. Third, it highlights similar heteroplasmy dynamics in *C. elegans*, mammalian tissues and human patients (Monnot et al., 2011, Kang et al., 2016). Our results are also consistent with age-dependent heteroplasmy shifts in mice, suggesting that the underlying mechanisms may be conserved (Jenuth et al., 1997).

Our results are complementary to recent studies measuring *C. elegans* heteroplasmy (Ahier et al., 2018, Ahier et al., 2021). While we isolate cells, the Zuryn lab developed a method to isolate mitochondria. The two methods provide parallel approaches to study tissue-specific mtDNA dynamics. For example, a limitation of our approach is that long neuronal projections are likely to be lost during cell isolation. Consequently, our data likely reflect mutant load in neural soma. Indeed, this may be a factor that explains why we observe an increase in mutant mtDNA levels, whereas the Zuryn’s group reports the opposite (Ahier et al., 2018). On the other hand, the cellular context and mitochondrial integrity may be compromised during mitochondrial isolation. In addition, our approach allows a measure of mutant mtDNA load in a cell-type specific manner across different conditions and time-points. Collectively, the two approaches are complementary strategies that provide fundamental insights into the mechanisms that regulate mtDNA dynamics in different tissues of *C. elegans*.

In addition to assaying heteroplasmy levels in muscles and neurons, we assessed the mechanisms behind the tissue-specific differences. Our results demonstrate that AMPK affects heteroplasmy and contributes to the differences between muscles and neurons. AMPK is a master regulator of cellular metabolism, which is activated in response to increased energy demands and metabolic stress (Mihaylova and Shaw, 2011). When activated, AMPK triggers multiple downstream targets to restore cellular homeostasis (Herzig and Shaw, 2018). Among these targets are key processes mediating mitochondrial turnover, including mitochondrial degradation (mitophagy) and biogenesis (Mihaylova and Shaw, 2011). Mitophagy eliminates non-functional mitochondria, while biogenesis replaces the loss of mitochondrial mass and mtDNA. AMPK helps maintain the balance between these two antagonistic processes, which is vital for proper mitochondrial function (Palikaras and Tavernarakis, 2014). However, the role of AMPK in modulating heteroplasmy levels was previously unexplored. Our data show that AMPK plays an important role in regulating mutant-to-wildtype ratio in muscles. This finding has important consequences, as AMPK plays a central role in regulating the metabolism of most mammalian cell types and may contribute to heteroplasmy differences between them (Mihaylova and Shaw, 2011).

The ratio changes in favor of the mutant in the absence of AMPK. This effect may be caused by a reduction in AMPK-mediated mitophagy. There are several reasons supporting this mechanism. First, mitophagy is a primary target of AMPK regulation (Egan et al., 2011). Second, mitophagy inhibition leads to increased heteroplasmy level in *C. elegans* (Gitschlag et al., 2016, Lin et al., 2016, Valenci et al., 2015, Ahier et al., 2021). It is possible that AMPK loss causes a tissue-specific decrease in mitophagy, leading to higher mutant load in muscles. Alternatively, the effect may be caused by a reduction in AMPK-mediated biogenesis. However, this possibility seems unlikely given that mitochondrial biogenesis favors mutant mtDNA proliferation (Lin et al., 2016, Gitschlag et al., 2020). Moreover, our data show that somatic mtDNA copy number declines continuously throughout *C. elegans* lifespan, suggesting more degradation than biogenesis. Therefore, we hypothesize that AMPK knockout effects are mediated primarily by mitophagy and not mitochondrial biogenesis.

In addition to heteroplasmy, we examined mtDNA copy number dynamics throughout development. We discovered that muscles, neurons, and other somatic cells lose the majority of their mtDNA content during development. Consistent with our findings, previous study reported a decline in somatic mtDNA throughout development (Tsang and Lemire, 2002a). Interestingly, our results show that the sharp drop in mtDNA levels is not accompanied by selection against mutant mtDNA. On the contrary, heteroplasmy level increases in neurons and remains stable in muscles as mtDNA levels decline. These findings are surprising, as we expected the mutant pathogenicity to increase due to mtDNA decline, leading to stronger selection against the mutant (Figure 7C) (Gitschlag et al., 2016, Lin et al., 2016, Valenci et al., 2015, Ahier et al., 2021). The lack of negative selection suggests that the age-related mtDNA decline does not preferentially target mutant mtDNA, at least in the heteroplasmic strains analyzed. We further examined whether the decrease in mtDNA copy number continues after development is complete. The data from mature adult animals (Day 8) show that mtDNA copy number keeps decreasing past the onset of adulthood (Figure 6D). By this point in maturity, somatic cells lose more than 80% of their starting mtDNA content. This age-related decline in mtDNA content also occurs in mammals, although at a much slower pace than what we observe in worms (Mengel-From et al., 2014). Given the scale of mtDNA loss, the drop in mtDNA copy number may be one of the key factors restricting *C. elegans* longevity. The speed of mtDNA loss suggests that it may be an actively regulated process.

The data from mammals indicates that mtDNA depletion has a detrimental effect on mitochondrial function (Coskun et al., 2012, Rice et al., 2014, Filograna et al., 2021). Our discovery raises the important question of how worms survive acute mtDNA loss at such scale. We hypothesize that the effects of mtDNA loss are mitigated by unique features of *C. elegans* and mitochondrial biology. First, while *C. elegans* have a short lifespan of 12-18 days on average (Johnson and Hutchinson, 1993), the electron transport chain proteins can have a relatively long half-life ranging from 4 to 46 days (Karunadharma et al., 2015). The combination of long protein half-life and short worm lifespan may reduce the need for continuous protein production by mtDNA. Second, flux control studies demonstrate that electron transport chain proteins are produced in large excess, which may further mitigate the effects of decreased protein production (Letellier et al., 1998). Overall, the factors described above may help *C*.*elegans* tolerate the decline in mtDNA content. On the other hand, they may allow worms to save energy and resources by limiting their investment in costly mtDNA maintenance mechanisms. In this case, somatic mtDNA decline may be advantageous given the unique features of *C*.*elegans* biology.

In this study, we have developed a *C. elegans* model that allows us to simultaneously examine three key variables affecting mtDNA mutations phenotype. This new method was successfully applied to muscles and neurons, giving new insights into heteroplasmy and copy number dynamics in different tissues. We have also identified a mechanism that contributes to heteroplasmy level differences between muscles and neurons. AMPK plays an important role in regulating mutant-to-wildtype ratio only in muscles. Our results uncover similar heteroplasmy dynamics between *C. elegans* and mammalian tissues, specifically, the fact that tissue-to-tissue differences emerge with age and are not present at early developmental stages. On the other hand, our studies bring to light striking differences in copy number dynamics between *C. elegans* and mammals. *C. elegans* somatic tissues go through a sharp mtDNA decline during development, losing half of their mtDNA by the first day of adulthood. mtDNA depletion at such a scale would presumably be detrimental for mammalian tissues. However, this loss does not cause drastic consequences in worms, as they continue to function throughout adulthood. This finding suggests that the importance of mtDNA maintenance may differ between short-lived and long-lived organisms, as well as different cell types. Our findings suggest that cell lifespan may be an important factor affecting the response of mammalian tissues to mtDNA mutations.

## Acknowledgements

We would like to thank Lantana Grub, James Held, Claudia Pereira and Kendal Broadie for the invaluable feedback on the manuscript. Some strains were provided by the CGC, which is funded by NIH Office of Research Infrastructure Programs (P40 OD010440). We thank David Miller’s Lab for strains and help with the adaptation of the cell isolation protocol for our needs. This work was generously supported by R01 GM123260 (MRP), the Discovery Award (PR170792) from the Department of Defense’s Congressionally Directed Medical Research Program (MRP), and Vanderbilt’s Trans-Institutional Program in Single Cell Biology (MRP). The VMC Flow Cytometry Shared Resource is supported by the Vanderbilt Ingram Cancer Center (P30 CA68485) and the Vanderbilt Digestive Disease Research Center (DK058404). Droplet Digital PCR to quantify transcript levels was performed through the Vanderbilt University Medical Center’s Immunogenomics, Microbial Genetics and Single Cell Technologies core.

## Author Contributions

Conceptualization, N.T. and M.R.P.; Methodology, N.T.; Validation, N.T.; Formal Analysis, N.T.; Investigation, N.T.; Resources, M.R.P.; Writing – Original Draft, N.T.; Writing – Review & Editing, N.T. and M.R.P.; Visualization, N.T.; Supervision, M.R.P.; Project Administration, M.R.P.; Funding Acquisition, M.R.P.

## Declaration of interests

The authors declare no competing interests.

## STAR Methods

### RESOURCE AVAILABILITY

There are three subheadings required in this section: lead contact, materials availability, and data and code availability. Lead contact

## Lead contact

Further information and requests for resources and reagents should be directed to and will be fulfilled by the lead contact, Maulik Patel

## Materials availability

New strains and reagents generated during this project are available through the lead contact.

## Data and code availability

The data reported in this paper will be shared by the lead contact upon request

No original code was generated during this study

### EXPERIMENTAL MODEL AND SUBJECT DETAILS

#### *C.elegans* culture conditions

*C*.*elegans* strains used in this study were cultured in two different sets of conditions depending on experimental stage. Prior to age synchronization, temperature-sensitive strains were kept at 16°C on standard nematode growth medium (NGM) plates seeded with OP50 *E*.*coli* bacteria. After age synchronization, half of L1 larvae population was used for cell isolation, while the other half was replated on 150mm 8P plates seeded with NA22 *E*.*coli* bacteria. The latter animals were incubated at 25°C until they reached the required developmental stage. Only hermaphrodite worms were used in the experiments. The experiments were performed with worms at the following developmental stages: L1 larva, Day 1 adult and Day 8 adult, as indicated. Detailed information on the strains used is included in the Key Resources table.

## METHODS DETAILS

### Genetic crosses

Classical genetic crosses were used to introduce *glp-1(e2141ts)* allele along with *zcIs21* or *evIs111* gene constructs to mutant mtDNA-carrying animals. The genotypes of the two gene constructs are provided in Key Resources table. *glp-1(e2141ts)* allele was added to ensure that worm population remains age-synchronized after reaching adulthood. It inhibits germline formation and, consequently, worm proliferation when animals are maintained at the restrictive temperature of 25°C. The two gene constructs, zcIs21 and *evIs111*, were used to induce tissue-specific GFP expression in body wall muscles and neurons, respectively. GFP-labeling was required to isolate the tissues of interest during cell sorting. *aak-2(ok524)* allele was introduced to *uaDf5*-carrying animals to test the effects of AMPK knockout on heteroplasmy level. The allele contains a deletion in *aak-2*, the catalytic subunit of AMPK kinase complex.

### Preparing worm population for embryo isolation

During this step worms were maintained at 16°C. First, they were grown on 90mm NGM plates seeded with OP50 *E*.*coli* bacteria. Once the food supply was exhausted worms were replated on 150mm P-plates seeded with NA22 *E*.*coli* bacteria. These plates allow us to grow sufficiently large worm populations for subsequent cell isolation. The P-plates were maintained at 16°C for 4-5 days until worms were just about to starve. This step was followed by embryo isolation.

### Embryo isolation

On the 5^th^ day after plating worms reached sufficient density and were washed off the plates with Milli-Q water into 50 ml conical tubes. Each tube should have enough volume for worms from 3-4 P-plates. The tubes were spun for 3 minutes at 1000xg. Supernatant was aspirated and 10 ml of bleach solution was added to each 50ml tube. The bleach solution was freshly made prior to embryo isolation, according to the recipe indicated below. The tubes were vortexed at low-speed mode and incubated on a rocking nutator for 5 minutes. All procedures were performed at room temperature. After incubation the reaction was stopped the addition of egg buffer to the total volume of 50 ml. The tubes were vortexed and spun for 3 minutes at 1000xg. The washing steps were repeated 3 more times. 50 ml tubes were switched to 15 ml conical tubes for the final 3^nd^ wash. These tubes are more convenient for embryo isolation. After the washes cells were resuspended in 10 ml of 30% sucrose solution and centrifuged for 3 minutes at 500xg. This step separates embryos from debris. Embryos floating on the top of the solution were aspirated with a pasteur pipette and transferred to a fresh 15-ml conical tube. No more than 5 ml of sucrose solution was transferred along with the embryos. To pellet the embryos Milli-Q water was added to the tubes to the total volume of 15 ml. The tubes were vortexed and centrifuged for 3 minutes at 3000xg. After that, embryos were resuspended in 25 ml of M9 solution and transferred to fresh 50ml tubes. The tubes were placed on a rocking nutator and incubated at room temperature overnight.

### Cell isolation

After overnight culture in M9 buffer, L1 larva hatch from the isolated embryos. To pellet down the worms, tubes were centrifugated at 1000g for 3 minutes. Then pellet was washed with MilliQ water, resuspended, and divided on two parts: one part for cell isolation from L1 animals, another part for cell isolation at later developmental stages. To isolate cells worms were transferred to 1.5 ml tube and spun at 14,000 rpm for 1 minute. Supernatant was removed and pellet was resuspended in SDS-DTT. SDS-DTT was added in 2-to-1 ratio to the pellet’s volume. The tubes were incubated on a rocking nutator for 2 minutes (L1 larvae) or for 4 minutes (adult animals) depending on worms’ developmental stage. The reaction was stopped by adding 1X egg buffer to total volume of 1.5 ml. The tubes were centrifugated at 14,000 rpm for 1 minute. Pellet was washed with egg buffer 5 more times and resuspended in protease solution (15mg/ml in egg buffer). Protease was added in 2-to-1 ratio to the volume of the pellet. The tubes were placed on a rocking nutator and incubated at room temperature for 20 minutes. During the incubation pellet was pipetted up and down 160 times with P-200 pipette to promote cell separation. The reaction was stopped by adding 1X egg buffer to total volume of 1.5 ml. Tubes were spun at 500 rcf for 3 minutes at 4 °C. Pellet was washed one more time, resuspended in 0.5 ml of cold 1X sterile egg buffer and spun at 100 rcf for 3 minutes at 4 °C. This step pellets down large debris, while the cells remain in the supernatant. Supernatant was aspirated and pipetted through a 35 μm filter-top on a 5ml falcon tube with a cell strainer cap. The tubes were spun at 100g for 1 minute and placed on ice until the cell sorting.

### Growth of age-synchronized worm population

To isolate cells from worms at Day 1 and Day 8 adult stages, L1 larvae were plated on 150mm P-plates seeded with NA22 *E*.*coli* bacteria. The plates were maintained at the restrictive temperature of 25°C, which inhibits germline development in *glp-1(e2141ts)* animals. After they reached the required developmental stage the worms were washed off the plates and used in cell isolation experiments.

### Cell sorting with FACS

Three controls were used to calibrate FACS machine before the experiment: GFP positive, DAPI positive and GFP/DAPI negative. Right before cell sorting DAPI viability dye was added to the samples to final concentration of 0.125 μg/ml. Viable cells from GFP positive and Total cell populations were sorted into separate 96-well plates (200 cells per well). Each well contained 10 μl of 0.5X lysis buffer with 0.1 mg/ml proteinase K. After sorting, the plates were sealed, spun for 1 minute at 200g and put on lysis in a thermocycler under following conditions: 1 hour at 60°C, then 15 minutes at 95°C. After lysis the plates were stored at -20 until mtDNA quantification was performed.

### Measuring mutant and wild-type mtDNA levels

Heteroplasmy level and mtDNA copy number were quantified with droplet digital PCR (ddPCR). First, template-primer mix was prepared by combining 8 μl of cell lysate with 2 μl of primer mix. After that 8 μl of template-primer mix were transferred to Eppendorf twin.tec™ 96-well semi-skirted plates, combined with 4.5 μl nuclease-free water and 12.5 μl EvaGreen supermix. The plates were sealed and centrifuged at 200g for 1 minute. Next, the plates were loaded into Bio-Rad Droplet Generator and processed using standard droplet generation program for EvaGreen samples. PCR was performed with annealing temperature of 58°C for uaDf5 primers and 57.5°C for *mptDf2* primers. After PCR the plates were loaded into Bio-Rad QX200 Droplet Reader and the number of mutant and wild-type mtDNA-carrying droplets was quantified with QuantaSoft software. Primers used for measuring *mptDf2* level: NT29, NT31, NT33, NT34. Primers used for measuring *uaDf5* level: MP433, MP434, MP435, MP436.

### QUANTIFICATION AND STATISTICAL ANALYSIS

Unless otherwise noted, unpaired t-test with Welch’s correction was used to compare experimental groups. The sample sizes are indicated in the figure legends. Sample size N represents the number of independent 200-cell batches used in the experiment. Statistical analysis was performed using Prism 8.0.2 software. All experiments were repeated at least 2 times.

### Data transformation

In this study the cell types of interest were labeled using tissue-specific GFP expression. As a result, every tissue was derived from a separate *C*.*elegans* strain. Since heteroplasmy level can differ significantly between the strains, we could not make direct comparison between the tissues. The tissue-to-tissue differences observed after direct comparison could be caused by mutant load differences between the strains. To account for this possibility, the heteroplasmy level of each tissue was normalized to its corresponding cell suspension control. Data transformation was performed according to the following formula: Δh = ln ((h (h0 – 1)) / (h0 (h – 1))), where h and h0 represent the heteroplasmy levels of the tissue and cell suspension control, respectively (Burgstaller et al., 2014, Johnston and Jones, 2016, Latorre-Pellicer et al., 2019). The transformation accounts for potential mutant load difference between the strains and allows us to compare heteroplasmy level across the tissue types and conditions.

## KEY RESOURCES TABLE

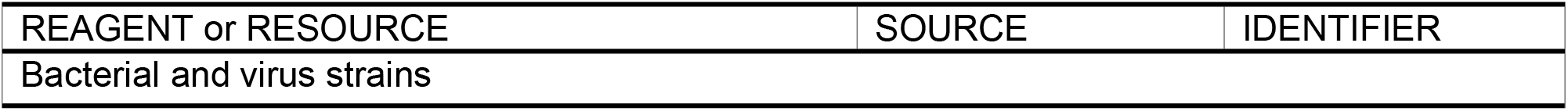
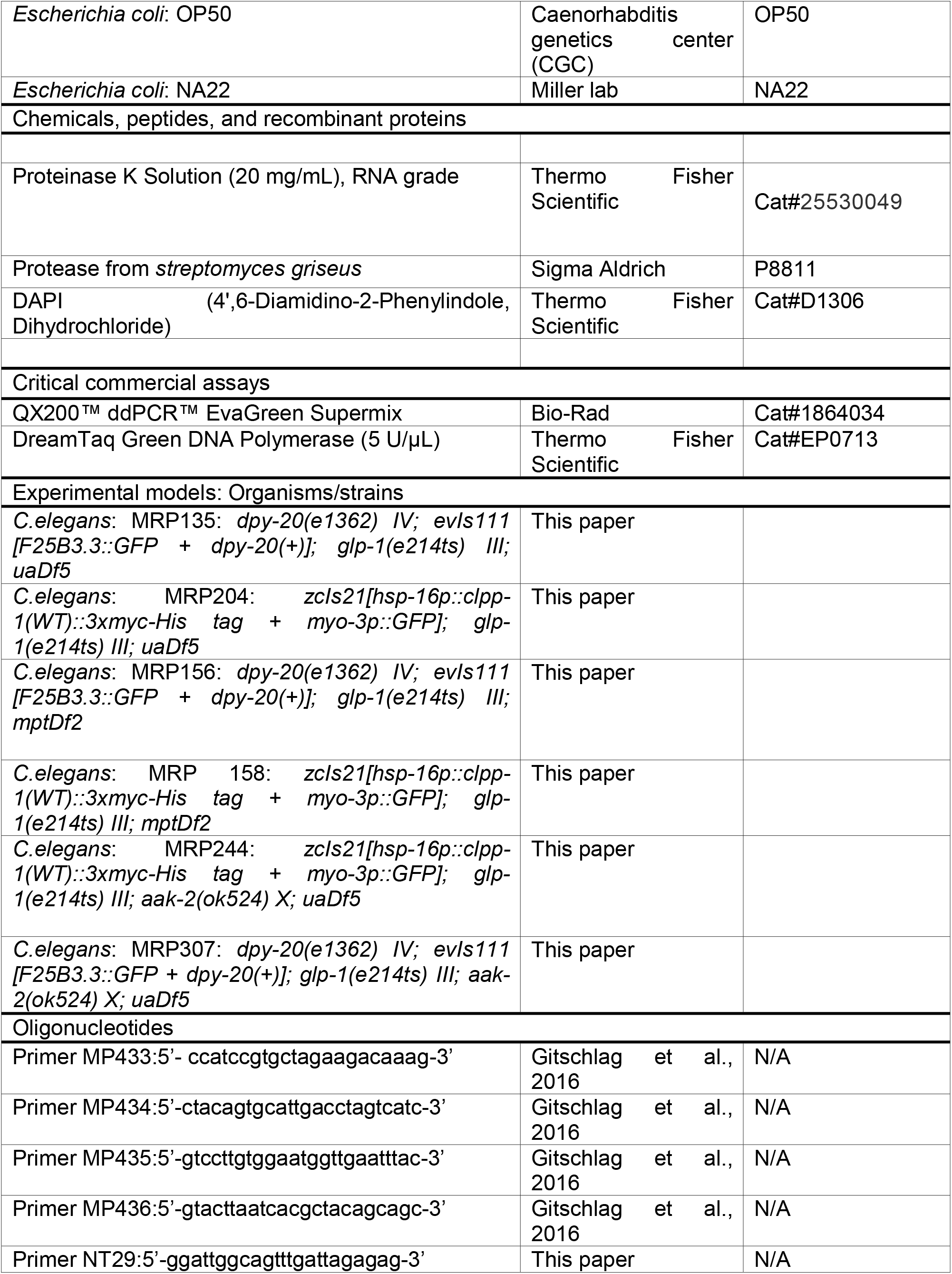
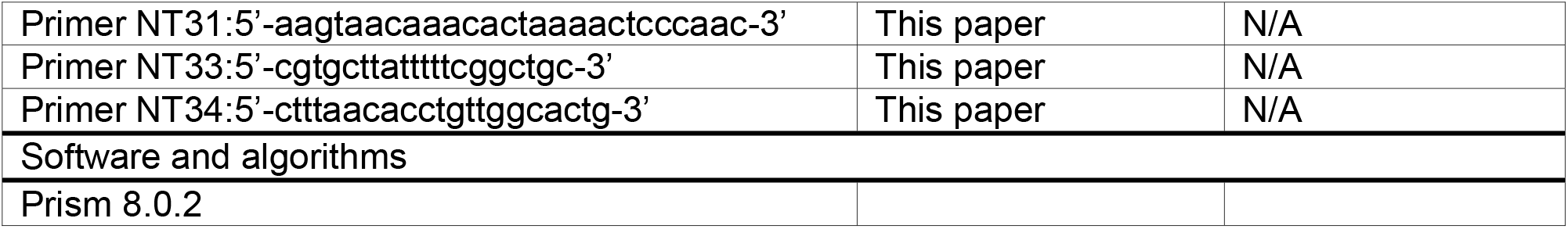

## Notes

### Competing Interest Statement

The authors have declared no competing interest.

